# Light power density governs local and systemic responses to photodynamic therapy

**DOI:** 10.64898/2026.11.10.765809

**Authors:** Hanqing Jiang, Eisuke Kanao, Yiming Xia, Stephanie Wagner, Steffen Hackbarth, Takahiro Nomoto

## Abstract

Photodynamic therapy (PDT) is a promising remedy against tumor cells by light activation of a photosensitizer, and its systemic antitumor effects are commonly attributed to immunogenic cell death and the release of soluble damage-associated molecular patterns (DAMPs). Here, we show that light power density (mW/cm^2^), which is often treated as a mere technical parameter, determines whether PDT suppresses distant, unirradiated tumors. At an identical light dose (J/cm^2^), PDT at a low power density sustained singlet oxygen (^1^O_2_) generation in tumors, preserved tumor perfusion, and suppressed both irradiated and unirradiated tumors. PDT at a high power density resulted in reduced ^1^O_2_ generation, reduced tumor perfusion, and increased programmed death-ligand 1 (PD-L1) expression in unirradiated tumors without inhibiting their growth. Serum levels of soluble DAMPs did not differ between the two conditions, whereas circulating small extracellular vesicles (sEVs) carried distinct protein cargo. Power density is thus a controllable determinant of the systemic effects of PDT, and circulating sEVs are candidate mediators of these effects.

**Teaser:** At an identical light dose, a lower power density extends the effect of photodynamic therapy beyond the illuminated tumor.

## INTRODUCTION

Light can be delivered with precise spatial and temporal control and has been used in numerous applications in chemistry, physics, biology, and medicine. Photodynamic therapy (PDT) uses this property to activate a drug only within the irradiated region and is clinically approved for the treatment of malignant tumors such as gliomas, esophageal cancers, and lung cancers (*1*). PDT consists of three components: light, a photosensitizer (PS), and oxygen (O_2_) (*2, 3*). Light excites the PS, which transfers energy to O_2_ and produces highly reactive singlet oxygen (^1^O_2_) (*4*), which is considered the main cytotoxic species. Beyond the direct cytotoxic effect, PDT also damages the vasculature and sometimes triggers rapid vascular shutdown (*5*) and subsequent hypoxia-induced tumor cell death (*6*), both of which can modulate the tumor microenvironment, inflammatory signaling (*7*), and immune cell infiltration (*8*). The mode of cell death is thought to determine these microenvironmental changes (*9*). Soluble damage-associated molecular patterns (DAMPs) released from dying tumor cells are widely regarded as hallmarks of PDT-induced immunogenic cell death (*10–12*). These molecules are considered to link PDT-induced cell death to systemic antitumor immunity (*13*), including the abscopal effect (*14*) (**Fig. 1A**). Because its photochemical mechanism differs from that of other therapeutics and its systemic toxicity is limited, PDT has been regarded as a promising partner for immunotherapy, and such combinations have shown efficacy in preclinical models (*15, 16*).

**Fig. 1.**
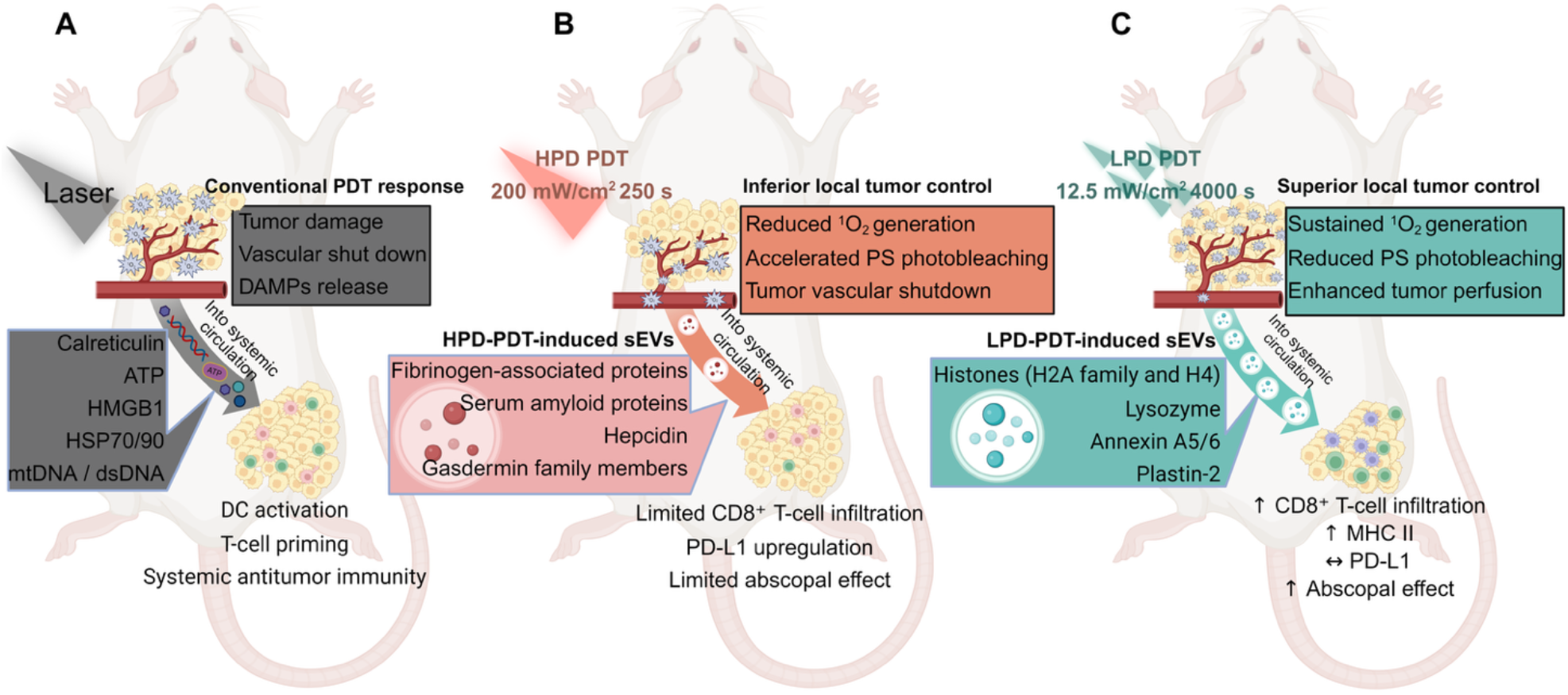
Conventional and power-density-dependent modes of action of PDT. **(A)** Conventional model of PDT-induced systemic antitumor immunity. PDT-mediated cell death promotes the release of DAMPs, followed by their entry into the systemic circulation. DAMPs induce activation of dendritic cells, antigen presentation, T-cell priming, and systemic antitumor immunity, ultimately contributing to the abscopal effect. **(B)** PDT at a high power density (HPD-PDT; 200 mW/cm^2^ for 250 s). HPD-PDT depletes O_2_, reduces ^1^O_2_ generation, accelerates PS photobleaching, and reduces tumor perfusion, resulting in inferior local tumor control. ^1^O_2_ is predicted to form mainly near blood vessels. HPD-PDT also yields sEVs enriched in coagulation and acute-phase proteins, with limited CD8⁺ T-cell infiltration, negligible MHC II increase, PD-L1 upregulation, and a weak abscopal effect. **(C)** PDT at a low power density (LPD-PDT; 12.5 mW/cm² for 4000 s). LPD-PDT sustains ^1^O_2_ generation, reduces PS photobleaching, and preserves tumor perfusion, resulting in improved local tumor control. LPD-PDT induces sEVs with an immunostimulatory protein profile and is associated with increased CD8⁺ T-cell infiltration, increased MHC II expression, unchanged PD-L1 levels, and an enhanced abscopal effect. HPD- and LPD-PDT deliver the same total light dose (50 J/cm²). Created in BioRender. JIANG, H. (2027). https://BioRender.com/nvfj4l2

However, systemic immune activation after PDT has not been reproduced consistently in the clinic, and its extent and mechanism remain debated even in preclinical work (*17, 18*). Because PDT kills tumor cells under most conditions, this inconsistency is difficult to explain simply by the release of soluble DAMPs alone, which implies an additional mechanism. In this context, light power density (mW/cm^2^), a parameter, which varies widely across studies yet has rarely been treated as a biological variable, may explain part of this inconsistency. In clinical practice, PDT is generally prescribed according to a defined total light dose (J/cm^2^), while power density is often set high to shorten treatment duration. Most preclinical studies have likewise used high power densities. However, the effects of PDT depend on power density even at a fixed total light dose (*19–21*). Lower power densities have been associated with stronger antitumor effects than higher power densities (*22*). Power density also influences the local inflammatory response (*23*). The effect of power density on local tumor control has been attributed to the balance between O_2_ consumption by the photochemical reaction and O_2_ supply to the tumor (*24*). However, this balance has rarely been measured in tumors during treatment, and whether ^1^O_2_ kinetics during PDT are related to systemic immunity has not been examined.

Here, we demonstrate that power density determines whether PDT acts as a local intervention or engages systemic antitumor immunity. Combining *in situ* photochemical monitoring with proteomic profiling of circulating small extracellular vesicles (sEVs) in a murine colon tumor model, we show that PDT at a high power density, similar to that used clinically, is associated with reduced ^1^O_2_ generation, substantial photobleaching of the PS, and decreased perfusion in the irradiated tumor, alongside an immunosuppressive microenvironment in the contralateral unirradiated tumor. PDT at a low power density, in contrast, preserves tumor perfusion and sustains ^1^O_2_ generation, reshaping the immune microenvironment in the contralateral unirradiated tumor and promoting abscopal effects. Serum levels of soluble DAMPs did not differ between the two conditions, whereas PDT at the low power density altered the protein cargo of circulating sEVs, including histones and proteins that mediate vesicle uptake by antigen-presenting cells (APCs), and increased antigen-presenting macrophages and CD8⁺ T cells in the distant tumor. Power density is thus a simple parameter that links local photochemical reactions with systemic immunity through a mechanism independent of soluble DAMPs **(Fig. 1, B and C)**.

## RESULTS

### Lower power density enhances local and abscopal effects at an identical light dose

We used talaporfin sodium as the PS because it is clinically approved for cancer treatment and its photophysical properties are well characterized (*25*). We first compared the effects of five power densities on irradiated and contralateral unirradiated tumors at a constant light dose of 50 J/cm^2^. As illustrated in **Fig. 2A**, BALB/c mice bearing bilateral subcutaneous CT26 tumors received intravenous talaporfin sodium (10 mg/kg) when the primary tumor reached approximately 200 mm^3^, and the primary tumor was irradiated at 680 nm 4 h later (**Table 1**). Tumor growth inhibition depended on the power density (**Fig. 2, B and C**). Lower power densities produced stronger suppression of the irradiated tumor, and 12.5 mW/cm^2^ produced the most effective control without recurrence during the observation period (**Fig. 2B**). Power densities of 12.5 to 50 mW/cm^2^ also inhibited the growth of contralateral unirradiated tumors (**Fig. 2C**).

**Fig. 2.**
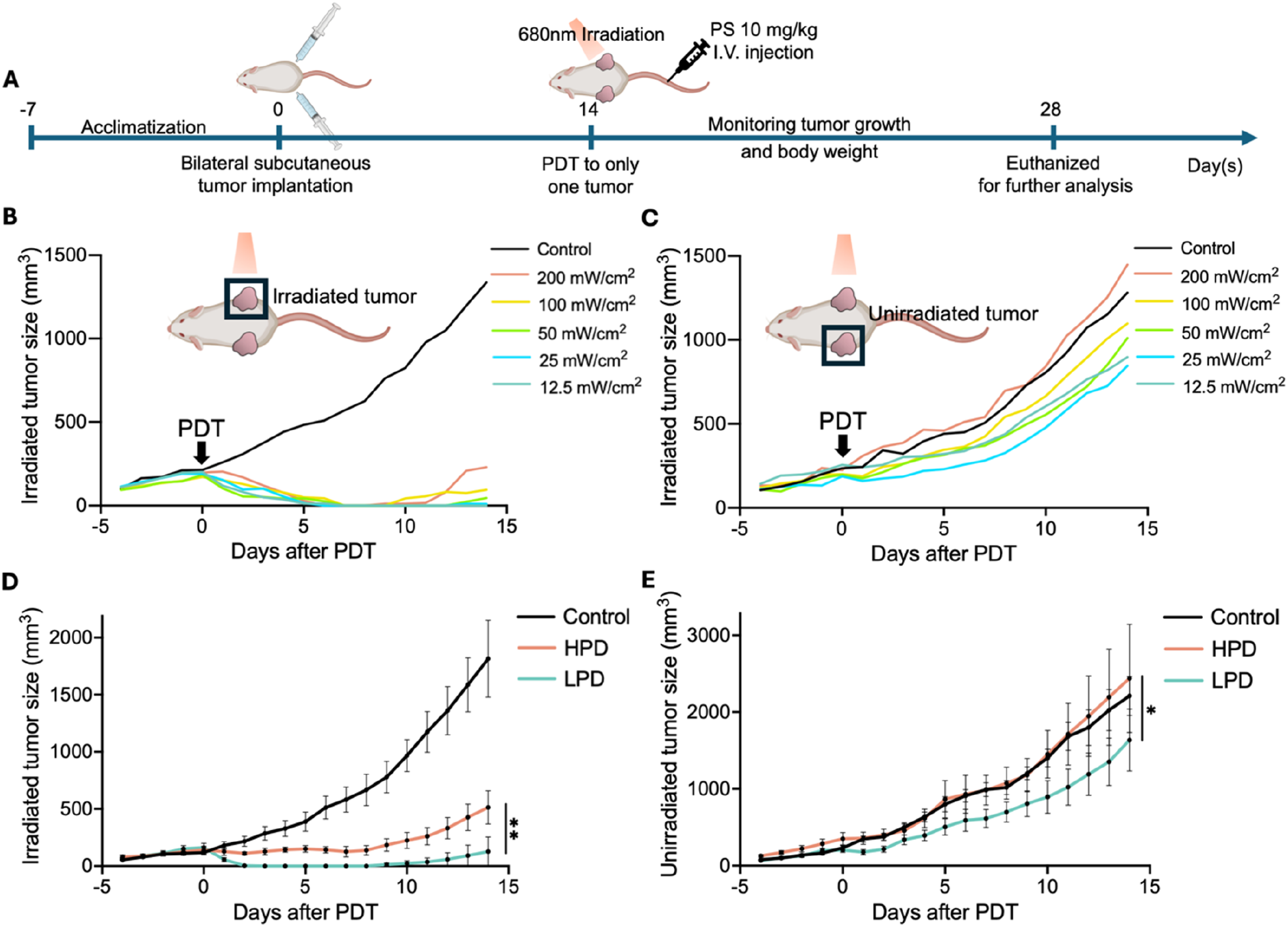
Lower power density enhances the local and abscopal effects of PDT at an identical light dose (50 J/cm^2^). **(A)** Schematic illustration of the experimental design. Mice bearing bilateral CT26 tumors received intravenous administration of PS, followed by PDT. Tumor growth was monitored for 14 days. **(B, C)** Growth curves of irradiated **(B)** and contralateral unirradiated **(C)** tumors in the preliminary experiment. The number of mice was n = 5 for the control and 100 mW/cm^2^ groups, and n = 4 for the other power-density groups. **(D, E)** Growth curves of irradiated **(D)** and contralateral unirradiated **(E)** tumors in an independent experiment comparing the control, LPD-PDT (12.5 mW/cm^2^), and HPD-PDT (200 mW/cm^2^) groups (n = 7 for control and LPD-PDT; n = 6 for HPD-PDT). Data are presented as mean ± SEM. Statistical analyses were performed using Tukey’s multiple comparisons test. *P < 0.05, **P < 0.01.

**Table 1.** Irradiation parameters used in this study. All irradiation conditions were delivered using an identical total light dose.

| Group | Irradiation power density (mW/cm<sup>2</sup>) | Irradiation time (s) | Irradiation dose (J/cm<sup>2</sup>) |
|---|---|---|---|
| Control | - | - | 0 |
| 1 | 200 | 250 | 50 |
| 2 | 100 | 500 |  |
| 3 | 50 | 1000 |  |
| 4 | 25 | 2000 |  |
| 5 | 12.5 | 4000 |  |

Based on these results, 12.5 mW/cm^2^ and 200 mW/cm^2^ were selected as representative low power density (LPD) and high power density (HPD) conditions, respectively, for the subsequent *in vivo* experiments. In an independent experiment, LPD-PDT suppressed both irradiated and contralateral unirradiated tumors more strongly than HPD-PDT (**Fig. 2, D and E**). Across the preliminary and main experiments, all 11 mice that survived until the end of the observation period showed complete tumor regression following LPD-PDT, with recurrence in only one mouse, yielding a tumor-free rate of 91% (10 of 11). In contrast, 5 of the 10 mice that survived until the end of the observation period in the HPD-PDT group showed complete tumor regression, and 4 of these 5 tumors recurred (**Table 2**). Thus, at an identical light dose, lowering the power density improved both local tumor control and the abscopal effect.

**Table 2.**
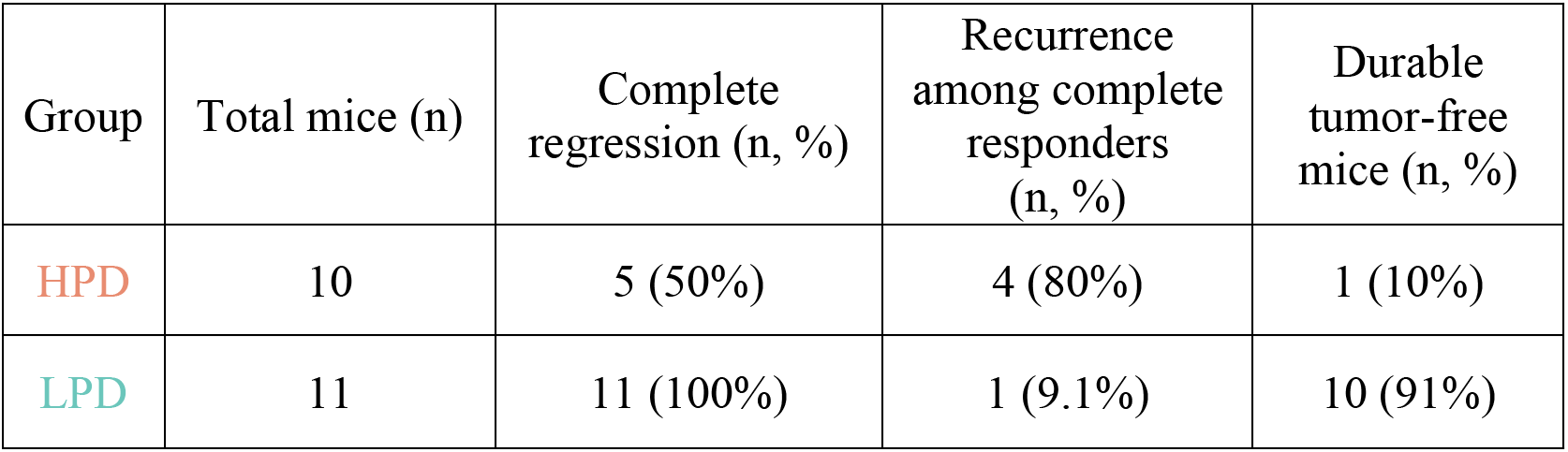
Therapeutic outcomes in the irradiated tumors. The results in Fig. 1, B and D are used. Complete regression is defined as the complete disappearance of the irradiated tumor during the observation period. Recurrence is defined as regrowth of a tumor after complete regression. Durable tumor-free survival is defined as complete tumor regression without subsequent recurrence during the observation period.

### High power density depletes O_2_ and reduces ^1^O_2_ generation in tumors

To examine how power density affects the photochemical reaction in tumors, we excited the PS with a custom-built nanosecond pulsed laser (∼400 ns) at 657 nm and detected ^1^O_2_ phosphorescence at 1270 nm in CT26 tumors *in vivo*. Pulsed excitation was required for time-resolved detection, whereas therapeutic PDT was performed using continuous-wave 680 nm irradiation; both wavelengths fall within the low-energy absorption band (Q band) of talaporfin sodium. In biomimetic environments such as aqueous albumin suspensions, this band is considerably broader than in phosphate-buffered saline (PBS) (*26*). Hence, 657 and 680 nm light are expected to excite the PS with comparable efficiency *in vivo* and to initiate the same photochemical pathway. Accordingly, we used the pulsed measurements, and the average power density was varied by changing the pulse repetition rate. To minimize inter-animal variability, all power densities were examined sequentially in the same tumor. Representative time-resolved ^1^O_2_ phosphorescence signals acquired from a tumor at the lowest (1.9 mW/cm^2^) and highest (120 mW/cm^2^) irradiation power densities, together with the corresponding fit curves, are shown in **Fig. 3A**. The decay curves were fitted using a biexponential model plus background to obtain the amplitude (*Amp*), reflecting the relative efficiency of ^1^O_2_ generation, and the characteristic decay parameters *τ*_1_ and *τ*_2_, corresponding to the apparent lifetimes of ^1^O_2_ and the triplet-state PS, respectively. In a previous study (*21*), it was shown that this *Amp* represents the balance between oxygen consumption and supply with lower values for increasing lack of oxygen. Three mice were examined; mice 1 and 2 were measured in descending order of power density, and mouse 3 in ascending order (**figs. S1 and S2**).

**Fig. 3.**
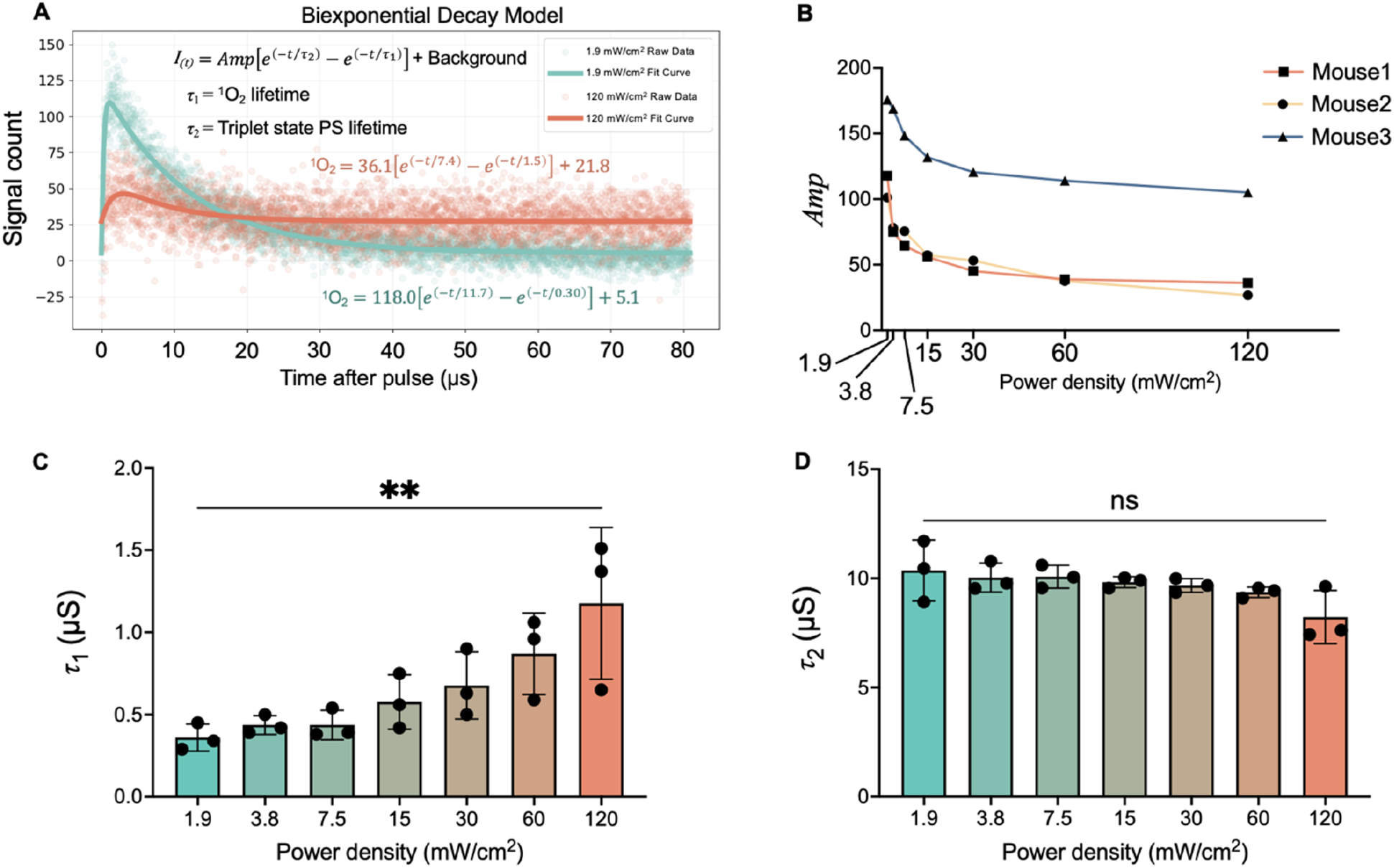
*In situ* monitoring of ^1^O_2_ photophysics within the irradiated tumor. **(A)** Representative time-resolved ^1^O_2_ phosphorescence curves acquired from one mouse at the highest (120 mW/cm^2^) and lowest (1.9 mW/cm^2^) power densities. Experimental data were fitted using a biexponential model. Different colors represent the highest and lowest irradiation power densities. Spheres, experimental data points; solid lines, fitted curves. **(B)** Amplitude (*Amp*) values obtained from biexponential fitting of ^1^O_2_ phosphorescence curves at seven power densities (1.9 to 120 mW/cm^2^). **(C)** Changes in the *τ*_₁_ parameter derived from biexponential fitting of ^1^O_2_ phosphorescence curves under seven different irradiation power densities. *τ*_₁_ represents the apparent lifetime of ^1^O_2_. **(D)** Changes in the *τ*₂ parameter derived from biexponential fitting of ^1^O_2_ phosphorescence curves under seven different irradiation power densities. *τ*_₂_ represents the apparent lifetime of the triplet-state PS. Data are presented as mean ± SD (n = 3 mice). Statistical analyses were performed using Tukey’s multiple comparisons test. \*\**P* < 0.01; ns, not significant.

Although the absolute *Amp* values differed between mice, all tumors exhibited the same inverse relationship between *Amp* and irradiation power density (**Fig. 3B**), indicating a higher ^1^O_2_-associated signal per excitation pulse at lower power densities. The *τ*_1_ value also increased at higher power densities (**Fig. 3C**). One possible explanation is that a certain fraction of the detected ^1^O_2_ signal may originate from intravascular regions, where ^1^O_2_ is subject to quenching by blood-borne molecules. If HPD irradiation also alters tumor blood perfusion, the replenishment of these quenching molecules may be reduced, allowing ^1^O_2_ to persist longer before encountering an effective quencher and thereby increasing the *τ*_1_. In contrast, preserved perfusion under LPD irradiation may facilitate continuous replenishment of blood-borne quenching molecules, resulting in more efficient quenching and a shorter *τ*_1_. This interpretation suggests that the dependence of *τ*_1_ on power density may reflect changes in the local quenching environment associated with tumor perfusion, rather than a change in the intrinsic lifetime of ^1^O_2_. In contrast, *τ*_2_, representing the apparent lifetime of the triplet-state PS, did not change substantially with power density (**Fig. 3D**). Together, these findings indicate that lower power density elicits a stronger ^1^O_2_-associated photodynamic response at an identical light dose, with higher *Amp* and shorter *τ*_1_, whereas *τ*_2_ remained largely unchanged. These photochemical findings are consistent with the superior control of irradiated tumor growth observed with LPD-PDT (**Fig. 2, B and D**).

### High power density accelerates PS photobleaching

To examine the effect of power density on PS photobleaching, we irradiated an aqueous PS solution (10 μM) with a halogen lamp at 100 or 25 mW/cm^2^ and recorded fluorescence spectra (600 to 800 nm), using fluorescence intensity as a measure of intact PS. Fluorescence decreased with increasing light dose under both conditions (**Fig. 4, A and B**), but the decrease was larger at 100 mW/cm^2^ than at 25 mW/cm^2^. Since the PS solution was exposed to the air and supplied with oxygen, these results suggest that the photobleaching can occur independently of O_2_ concentration to some extent. We next investigated photobleaching in the CT26 tumor (**Fig. 4C**). Tumors were irradiated with a 680 nm diode laser 4 h after PS administration, and PS fluorescence from the tumor was recorded during irradiation **(Fig. 4, D and E)**. After 7.5 J/cm^2^ at HPD (200 mW/cm^2^), tumor fluorescence decreased to approximately 40% of the pre-irradiation value (**Fig. 4F**), whereas it remained at approximately 50% after the same light dose at LPD (12.5 mW/cm^2^). Together, these results indicate that higher power density promotes PS photobleaching under both cell-free and tumor conditions. The greater photobleaching may contribute to the lower ^1^O_2_ generation observed during HPD-PDT (**Fig. 3B**), as photobleaching reduces the amount of intact PS available for further ^1^O_2_ generation. Photobleaching and oxygen limitation may thus act as interacting processes that restrict PDT activity at HPD.

**Fig. 4.**
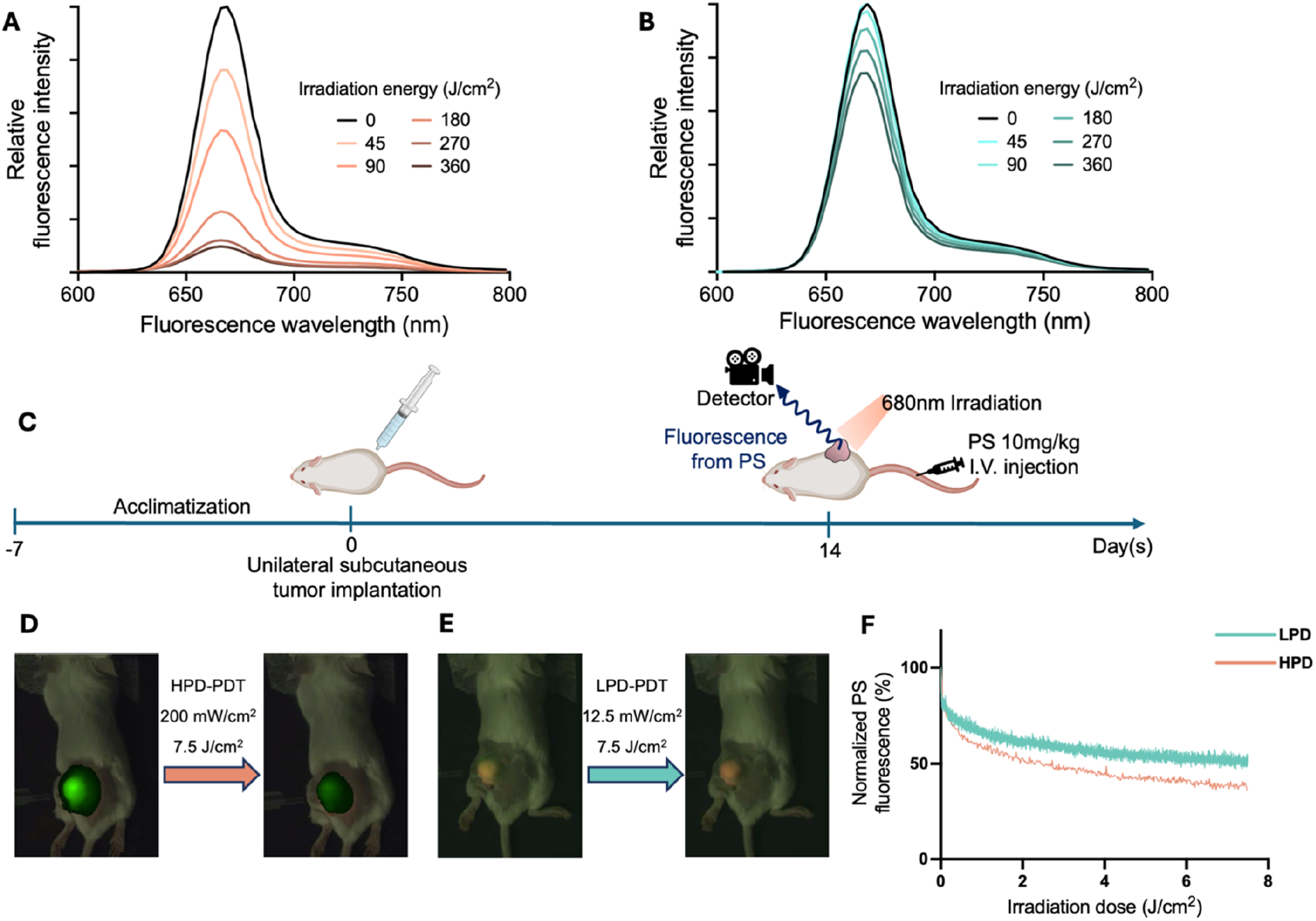
Photobleaching *in vitro* and *in vivo*. (A, **B)** Fluorescence spectra of talaporfin sodium (10 µM) in water. The samples were irradiated with a halogen lamp at 100 **(A)** and 25 **(B)** mW/cm^2^. **(C)** Schematic illustration of the *in vivo* experimental workflow for monitoring PS fluorescence during PDT. **(D, E)** Representative *in vivo* fluorescence images before and after irradiation. The tumors were irradiated under HPD **(D)** and LPD **(E)** conditions at 7.5 J/cm^2^. The fluorescence signals are shown in green. **(F)** Normalized PS fluorescence intensity in irradiated tumors as a function of delivered light dose under different irradiation power densities. Fluorescence signals were corrected using a non-irradiated reference region and normalized to the pre-irradiation value (100%). Data are presented as mean (n = 2 mice per group).

### High power density reduces tumor perfusion and increases serum IL-6

We measured tumor blood perfusion by laser speckle imaging (**Fig. 5A**). Before irradiation, perfusion in the tumor was comparable to that in normal tissue (**Fig. 5B**). HPD-PDT substantially reduced perfusion in the irradiated tumor (**Fig. 5C**). Because O_2_ is supplied by the blood, HPD-PDT may generate ^1^O_2_ mainly in and near the vessels (*27*), causing acute oxidative damage to the vessel’s wall, vascular shutdown, and reduced blood perfusion. In contrast, LPD-PDT slightly increased tumor perfusion (**Fig. 5, D and E**). At LPD, ^1^O_2_ is generated more slowly, which may not be sufficient to cause acute vascular damage and coagulation. In addition, ^1^O_2_ regulates vascular tone (*28, 29*), which may contribute to the upregulated perfusion. This finding is also consistent with our previous interpretation of the apparent ^1^O_2_ lifetime in **Fig. 3C**. The substantial reduction in blood perfusion after HPD-PDT slows the replenishment of blood-borne quenchers, thereby allowing ^1^O_2_ to persist longer before encountering an effective quencher and contributing to the observed increase in the apparent ^1^O_2_ lifetime (*τ*₁).

**Fig. 5.**
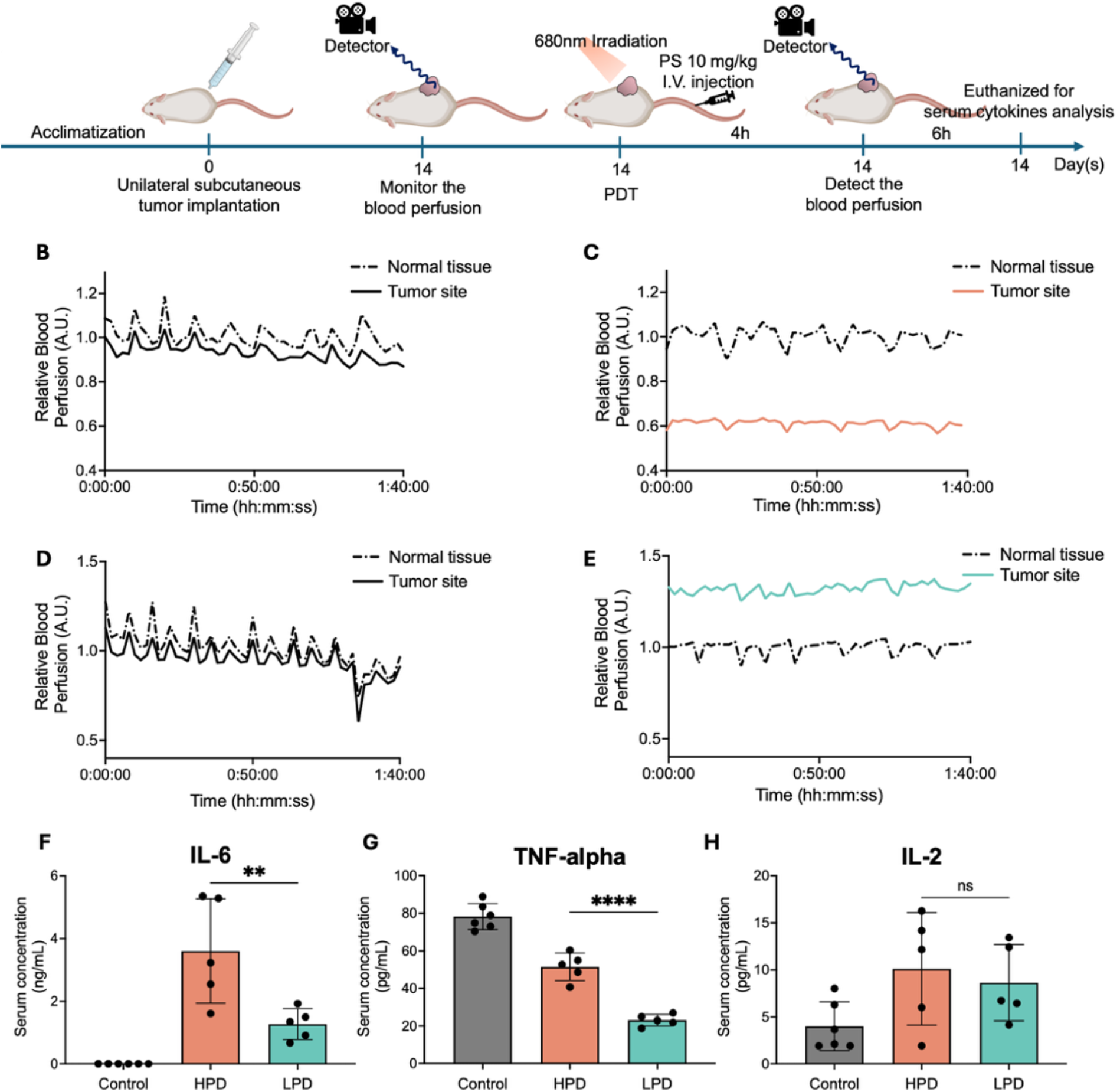
Blood perfusion and systemic inflammatory responses. **(A)** Schematic illustration of the experimental workflow. Tumor-bearing mice received PDT, followed by laser speckle pattern imaging and serum cytokine analysis. **(B–E)** Quantification of blood perfusion. The blood perfusion was monitored before **(B)** and after HPD-PDT **(C)** or before **(D)** and after LPD-PDT **(E)**. Skin distant from the tumor was defined as the normal tissue. **(F–H)** Serum levels of cytokines. IL-6 **(F)**, TNF-α **(G)**, and IL-2 **(H)** were quantified 6 h after PDT. Data are presented as mean ± SD. Sample sizes for the control, HPD-PDT, and LPD-PDT groups were n = 6, 5, and 6, respectively. Statistical analyses were performed using Tukey’s multiple comparisons test. \*\**P* < 0.01, \*\*\*\**P* < 0.0001; ns, not significant.

Serum cytokines also reflected the vascular response to HPD-PDT. Six hours after PDT, serum interleukin-6 (IL-6) was significantly higher in the HPD-PDT group than in the control and LPD-PDT groups (**Fig. 5F**). As IL-6 activates coagulation pathways during inflammatory responses (*30*), the elevated IL-6 may reflect a prothrombotic response to vascular damage after HPD-PDT. The smaller increase after LPD-PDT suggests a milder vascular effect, consistent with the perfusion data **(Fig. 5, B to E).** In contrast, serum tumor necrosis factor-α (TNF-α) was highest in the control group and decreased after PDT, with the lowest level observed in the LPD-PDT group **(Fig. 5G).** Assuming that circulating TNF-α in tumor-bearing mice reflects the systemic inflammatory burden of the tumor, its decrease after PDT suggests the direct oxidative damage to the tumor cells, and the larger decrease after LPD-PDT indicates stronger killing effects, which should be also in line with the results in the *in situ* monitoring of ^1^O_2_ (**Fig. 3**). These results agree with theoretical studies showing that the spatial distribution of O_2_ and therefore ^1^O_2_ depends on power density (*29, 31*). Under HPD conditions, ^1^O_2_ is predicted to be produced mainly within well-oxygenated tumor vessels, making endothelial cells the primary target, whereas under LPD conditions ^1^O_2_ extends into the often hypoxic tumor parenchyma. Meanwhile, serum interleukin-2 (IL-2) did not differ significantly among the three groups (**Fig. 5H**). Because IL-2 is produced mainly by activated T cells, this result suggests that systemic T cell responses had not developed at this early time point.

### Low power density activates antitumor immunity in unirradiated tumors

To investigate mechanisms underlying abscopal effects, we analyzed immune cells and cytokines in the unirradiated tumors 14 days after PDT (**Fig. 6A**). The proportion of F4/80^+^ tumor-associated macrophages (TAMs) did not differ among the groups (**Fig. 6B**). However, LPD-PDT increased the expression of major histocompatibility complex class II (MHC II) on TAMs (**Fig. 6C**), whereas the expression of CD206 (an M2 macrophage marker) did not differ among the groups (**Fig. 6D**). LPD-PDT might shift TAMs toward an antigen-presenting phenotype. The proportion of CD3^+^ T cells was comparable among the groups (**Fig. 6E**), but LPD-PDT increased the proportion of CD8^+^ cells within the CD3^+^ population (**Fig. 6F**), consistent with activation of antitumor immunity.

**Fig. 6.**
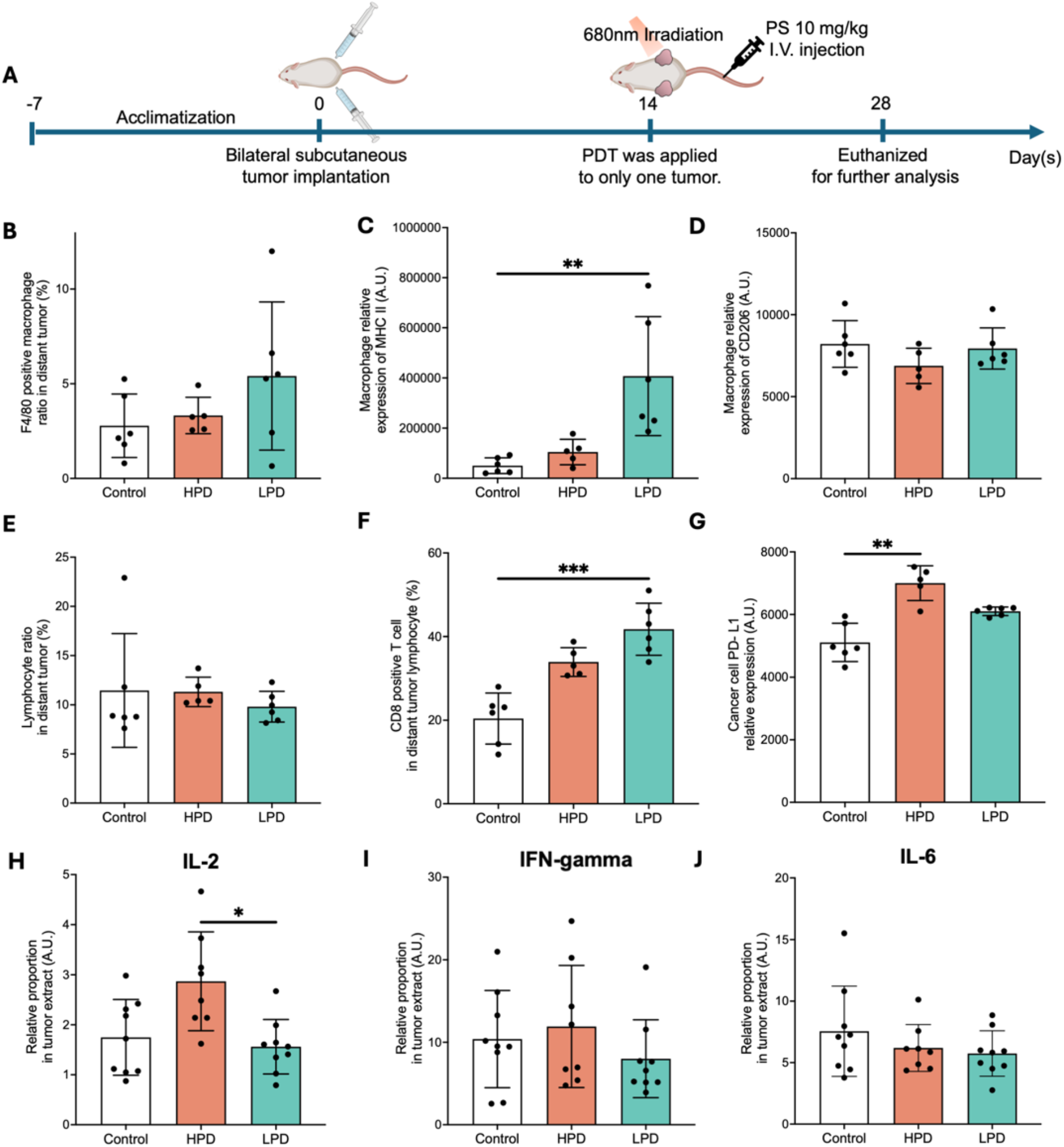
Immune microenvironment of unirradiated tumors. **(A)** Schematic illustration of the experimental workflow. Tumor-bearing mice received PDT, and immune profiling of the unirradiated tumors was performed 14 days after treatment. **(B)** Percentage of infiltrating macrophages (F4/80^+^ cells). **(C, D)** Expression levels of MHC II **(C)** and CD206 **(D)** on F4/80^+^ cells. **(E)** Percentage of infiltrating lymphocytes (CD3^+^ cells). **(F)** Percentage of CD8^+^ cells within the CD3^+^ population. **(G)** PD-L1 expression on tumor cells isolated from unirradiated tumors. **(H–J)** Levels of cytokines. IL-2 **(H)**, IFN-γ **(I)**, and IL-6 **(J)** in lysates of unirradiated tumors were quantified. Data are presented as mean ± SD. Sample sizes for the control, HPD-PDT, and LPD-PDT groups were n = 6, 5, and 6, respectively, for **(B–G)** and n = 9, 8, and 9, respectively, for **(H–J)**. Statistical analyses were performed using Tukey’s multiple comparisons test. \**P* < 0.05, \*\**P* < 0.01, \*\*\**P* < 0.001.

Interestingly, programmed death-ligand 1 (PD-L1) expression on tumor cells increased after HPD-PDT but not after LPD-PDT (**Fig. 6G**). The upregulation of PD-L1 suggests that HPD-PDT may induce a compensatory immunosuppressive response in unirradiated tumors. We also measured cytokines in lysates of unirradiated tumors by enzyme-linked immunosorbent assay (ELISA). IL-2 was elevated only in the HPD-PDT group (**Fig. 6H**), whereas interferon-γ (IFN-γ) and IL-6 did not differ among the groups (**Fig. 6, I and J**). The elevation of IL-2, together with increased PD-L1 expression but without an increase in CD8^+^ T cells or IFN-γ, suggests that the immune response induced by HPD-PDT did not lead to effective cytotoxic immunity. Because prolonged IL-2 signaling maintains regulatory T cells (*32*), the elevated IL-2 in the HPD-PDT group may also reflect an immunosuppressive milieu.

### Circulating soluble DAMPs

We next sought to determine whether circulating soluble DAMPs, which are widely recognized as key mediators linking local PDT-induced damage to systemic immune activation, were responsible for the differential immune responses induced by different power densities. Considering that immunogenic cell death and eventual soluble DAMP release are initiated shortly after PDT, serum samples were collected at 0.5, 6, and 24 h after treatment to quantify three representative DAMPs: high mobility group box 1 (HMGB1), heat shock protein 70 (HSP70), and adenosine triphosphate (ATP). Serum HMGB1 did not differ among the three groups at 0.5 h after PDT (**Fig. 7A**). At 6 h, both PDT-treated groups exhibited significantly lower levels than the control group, whereas no difference was observed between the HPD and LPD groups (**Fig. 7B**). Although HMGB1 is a well-characterized DAMP released during cellular stress and death, its circulating concentration reflects not only release from damaged tissues but also extracellular binding, cellular uptake, and clearance. Thus, the lower serum HMGB1 levels observed after PDT do not necessarily indicate reduced local HMGB1 release and may reflect the dynamic balance between release and systemic clearance at this early time point. At 24 h, HMGB1 was comparable in all groups (**Fig. 7C**). HSP70 did not differ among the groups at 0.5 h (**Fig. 7D**). At 6 h, HSP70 was higher in both PDT groups than in the control group, with no difference between HPD and LPD (**Fig. 7E**). At 24 h, the differences among the groups were not significant (**Fig. 7F**). Serum ATP increased rapidly and transiently after PDT; it was higher in both PDT groups than in the control group at 0.5 h (**Fig. 7G**) and remained significantly higher at 6 h, although the overall ATP levels declined relative to those at 0.5 h (**Fig. 7H**). At 24 h, ATP levels decreased to the baseline with no significant differences among groups (**Fig. 7I**). Thus, none of the three DAMPs differed between HPD-PDT and LPD-PDT at any time point, indicating that the stronger systemic response to LPD-PDT cannot be explained by the levels of these soluble DAMPs alone.

**Fig. 7.**
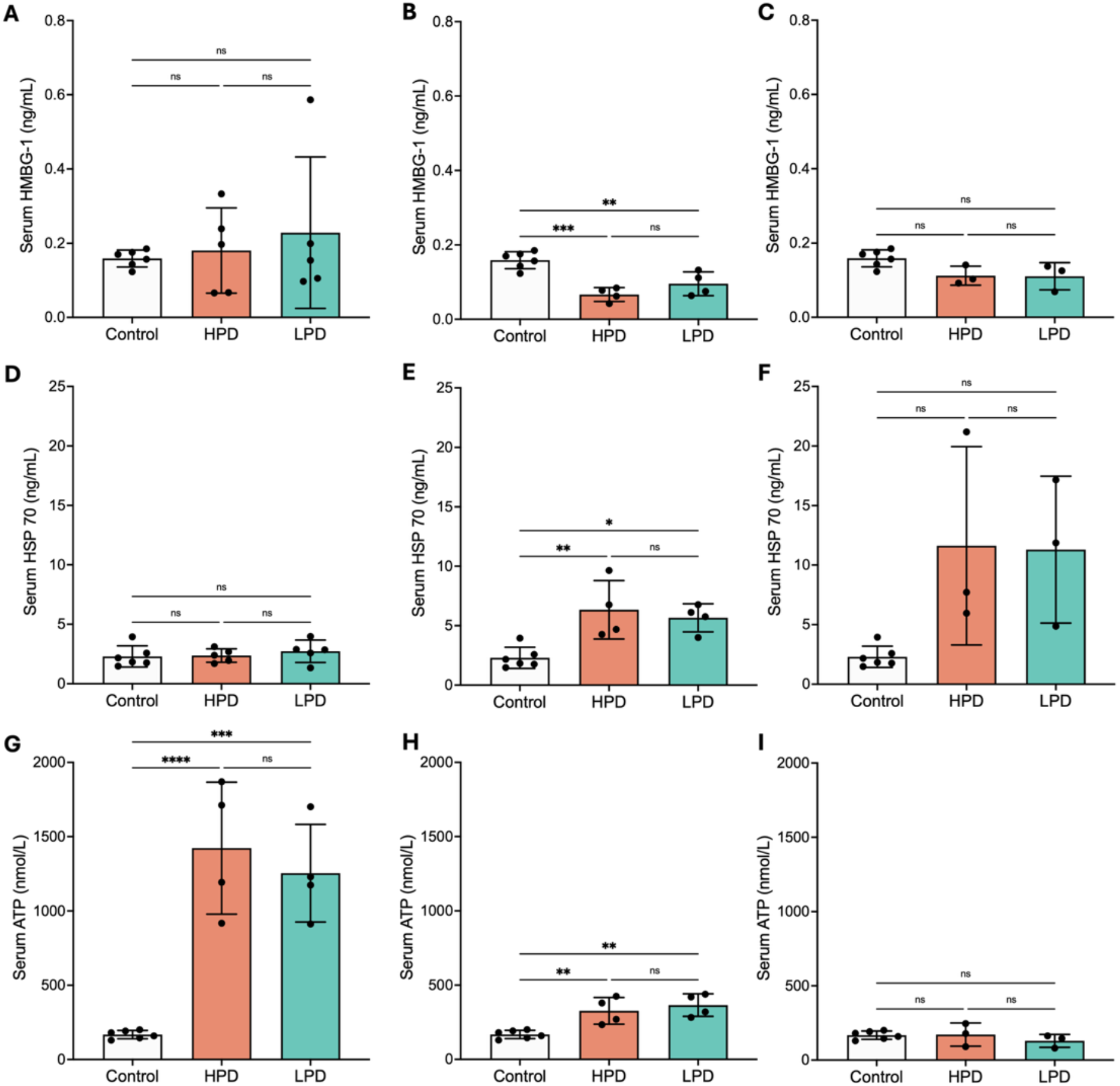
Temporal changes in circulating soluble DAMP levels following photodynamic therapy. **(A–C)** Serum HMGB1 concentrations measured at 0.5 h **(A)**, 6 h **(B),** and 24 h **(C)** after PDT. **(D–F)** Serum HSP70 concentrations measured at 0.5 h **(D),** 6 h **(E),** and 24 h **(F)** after PDT. **(G–I)** Serum ATP concentrations measured at 0.5 h **(G),** 6 h **(H),** and 24 h **(I)** after PDT. The same six control mice, which did not receive PDT, were used as the control group at all three time points. Sample sizes for the control, HPD-PDT, and LPD-PDT groups were n = 6, 5, and 5, respectively, at 0.5 h; n = 6, 4, and 4, respectively, at 6 h; and n = 6, 3, and 3, respectively, at 24 h. Data are presented as mean ± SD. Statistical analyses were performed using Tukey’s multiple comparisons test. \**P* < 0.05, \*\**P* < 0.01, \*\*\**P* < 0.001, \*\*\*\**P* < 0.0001; ns, not significant.

### Power density alters the protein cargo of circulating sEVs

To investigate whether PDT-treated tumor cells release macrophage-regulating signals independently of direct cell–cell contact, bone marrow-derived macrophages (BMDMs) were cultured in conditioned media (CM) from CT26 cells subjected to halogen-lamp irradiation at 100 or 20 mW/cm², with both conditions delivering a total light dose of 60 J/cm². Both HPD- and LPD-PDT-derived CM altered the expression of macrophage-associated markers **(fig. S3)**. LPD-PDT-derived CM significantly increased CD86 and TNF-α expression compared with both control and HPD-PDT-derived CM. CD206 expression was also significantly increased by both PDT-derived CM, with a greater increase following LPD-PDT. In contrast, Ym-1 and PPAR-γ expression decreased following both treatments, with no significant differences between HPD and LPD, while Irf-5 expression was significantly reduced only following LPD-PDT. These findings indicate that PDT-treated CT26 cells release macrophage-regulating mediators and that the resulting macrophage response varies with irradiation power density.

Because CM contain both soluble factors and extracellular vesicles (EVs), the contribution of each to the observed macrophage responses remains unresolved. Given the potential role of circulating small extracellular vesicles (sEVs) in tumor–immune communication and the possibility that PDT-induced vascular changes influence their release and systemic distribution, we hypothesized that irradiation power density alters the protein cargo of circulating sEVs. To test this, plasma sEVs were isolated 6 h after PDT and analyzed by liquid chromatography– tandem mass spectrometry (LC-MS/MS). The size distributions of the isolated sEVs are shown in **Fig. 8A**, with a modal diameter of approximately 100 nm in all groups. Spearman correlation analysis and hierarchical clustering revealed clustering by treatment group **(Fig. 8B)**, suggesting that PDT at different power densities was associated with distinct circulating sEV proteomic profiles. To investigate the molecular mechanisms governing power-density-dependent systemic responses, we compared the sEV proteomes between groups (LPD-PDT versus control, HPD-PDT versus control, and HPD-PDT versus LPD-PDT; **Fig. 8, C to E**; **table S1; data file S1**). Compared with the control and LPD-PDT groups, sEVs after HPD-PDT were enriched in proteins associated with the acute-phase response (*33, 34*), vascular damage, and coagulation (*35, 36*). Compared with the control, HPD-PDT increased the acute-phase proteins serum amyloid A1 (SAA1), SAA2, and C-reactive protein (CRP) (*33*), the myeloid activation marker CD14, the endothelial damage marker von Willebrand factor (vWF) (*35*), coagulation factors VII, XI, and XIII, and fibrinogen chains (*35, 36*), and gasdermin A2 (GSDMA2), a pore-forming protein involved in lytic cell death. This protein profile is consistent with the vascular damage and systemic inflammatory response observed after HPD-PDT (**Fig. 5**). HPD-PDT also decreased the endogenous anticoagulant tissue factor pathway inhibitor (TFPI) (*37*) and phosphatidylserine-binding proteins that mediate vesicle uptake by phagocytes, including lactadherin and annexins A5 and A6 (*38*), which are involved in the recognition and uptake of phosphatidylserine-exposing vesicles by phagocytes. The reduced abundance of these proteins may affect sEV uptake by phagocytes, potentially including APCs. These changes may be relevant to the unchanged MHC II expression observed in the contralateral unirradiated tumors after HPD-PDT (**Fig. 6C**), suggesting a potential association between altered circulating sEV composition and the antigen-presenting phenotype of the tumor microenvironment. In contrast, sEVs after LPD-PDT carried proteins associated with immunogenic cell death (*39, 40*). Compared with both the control and HPD-PDT groups, they were enriched in the DAMP-associated proteins histones H2A and H4 and lysozyme C-1 (*39, 40*). Proteins that mediate vesicle uptake by APCs, including annexins A5 and A6 (*38*), remained at control levels after LPD-PDT but decreased after HPD-PDT. LPD-PDT sEVs also contained more fatty acid-binding protein 5 (FABP5), a regulator of macrophage polarization (*41*), than sEVs from the other groups. This finding may be relevant to the altered macrophage phenotype observed in the contralateral unirradiated tumors after LPD-PDT (**Fig. 6C**). In addition, LPD-PDT-induced sEVs contained more plastin-2 (LCP1), an actin-bundling protein involved in immune synapse formation (*42*), than control sEVs. These results suggest that sEVs after LPD-PDT carry DAMP-like proteins such as histones, as well as macrophage-regulating proteins, while retaining the proteins that mediate their uptake by APCs.

**Fig. 8.**
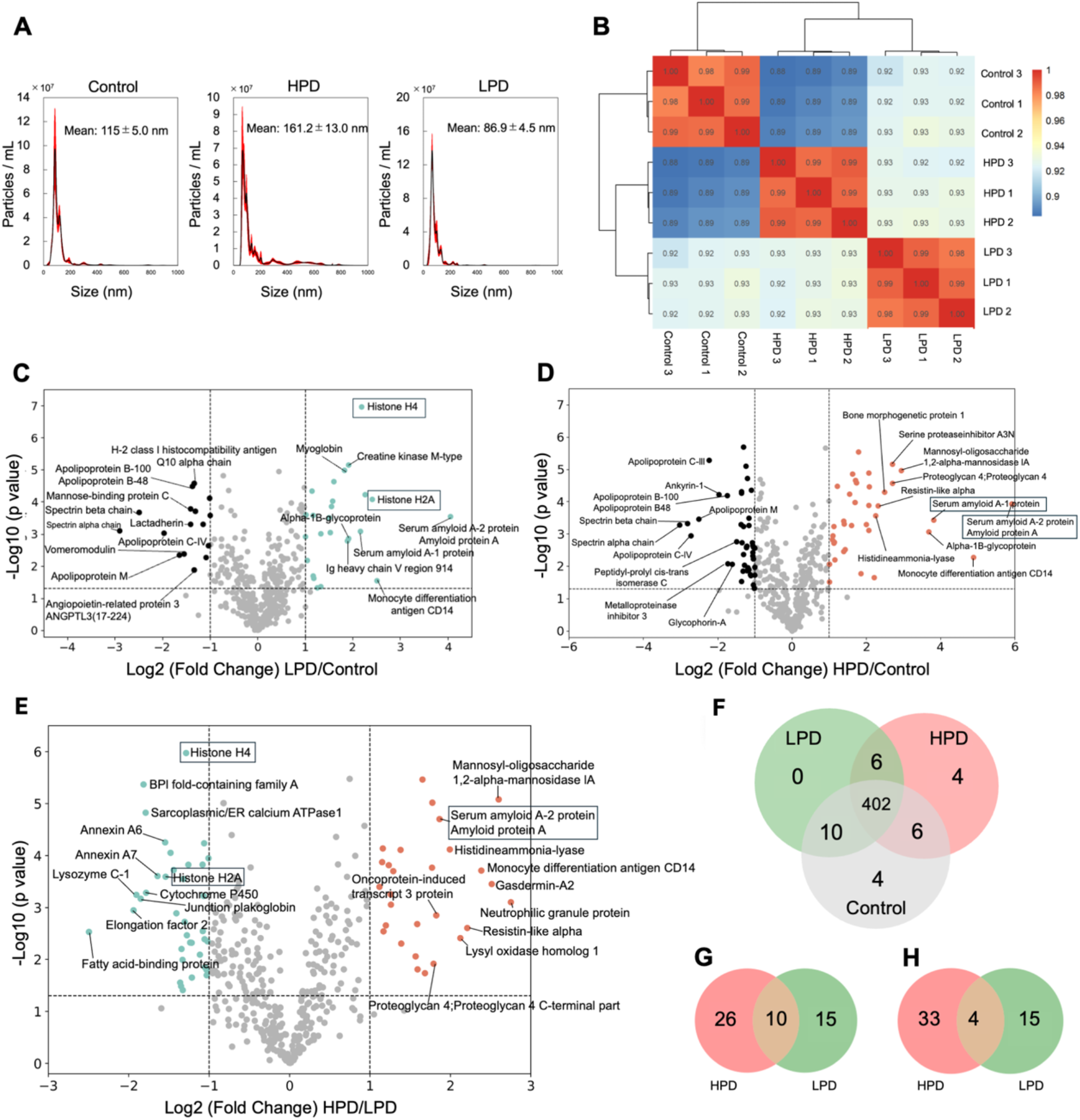
Irradiation power density shapes the circulating sEV proteome following PDT. **(A)** Size distribution of circulating sEVs isolated 6 h after PDT. **(B)** Spearman correlation heatmap with hierarchical clustering. **(C–E)** Volcano plots of LPD versus control **(C)**, HPD versus control **(D)**, and HPD versus LPD **(E)**; proteins with the largest fold changes are labeled. **(F–H)** Venn diagrams of proteins identified in each group **(F)** and of proteins significantly upregulated **(G)** or downregulated **(H)** in the HPD and LPD groups relative to the control group. Proteins quantified in at least two of three biological replicates in at least one group were analyzed, and those with fold change ≥ 2 and two-sided *P* < 0.05 were considered significant.

Most proteins were shared among the groups, with 402 proteins detected in all three groups (**Fig. 8F**). Relative to the control, HPD-PDT increased 36 proteins and decreased 37, whereas LPD-PDT increased 25 proteins and decreased 19 (**Fig. 8, G and H**). Only 10 increased proteins (**Fig. 8G**) and 4 decreased proteins (**Fig. 8F**) were shared between the two conditions, indicating that the two power densities altered different sets of proteins rather than the same proteins to different extents. These differences in sEV cargo may contribute to the macrophage activation and CD8⁺ T cell infiltration in unirradiated tumors after LPD-PDT.

## DISCUSSION

In this study, LPD-PDT and HPD-PDT used the same PS dose, wavelength, and total light dose (50 J/cm^2^) and differed only in the rate of light delivery. Under these conditions, LPD-PDT suppressed the growth of both the irradiated and the unirradiated tumors, whereas HPD-PDT was less effective against the irradiated tumor and did not affect the unirradiated one (**Fig. 2, D and E**). Thus, the generation or transmission of a signal acting outside the irradiated field depends on power density. Serum HMGB1, HSP70, and ATP were comparable between the two treatment groups at all the examined time points (**Fig. 7**), and the difference in the systemic effect is therefore unlikely to be explained by the amount of these soluble DAMPs. By contrast, the circulating sEVs had markedly different protein profiles (**Fig. 8**). The vascular responses also differed; tumor perfusion was substantially reduced by HPD-PDT, while it was maintained or slightly increased by LPD-PDT (**Fig. 5**). Because sEVs released in the tumor require blood or lymphatic flow to reach lymphoid tissues and the distant tumor, these results can be explained if the sEVs generated during the irradiation are exported through the tumor vasculature and if such export is restricted when the vasculature is shut down. We did not measure sEV transport directly, and this interpretation remains speculative. Nevertheless, it adds a step to the current model of PDT-induced immunity: the local photochemical reaction determines what is released, and perfusion during and after irradiation determines whether the released material leaves the tumor. Blood flow during PDT may thus be not only a target of treatment but also a route for the sEVs released during treatment. This interpretation is compatible with the concept of photodynamic priming, in which subcurative photodynamic treatment is used to modulate the tumor microenvironment rather than primarily to induce tumor cell killing (*43*). Mild photodynamic treatment increases local blood flow and tissue oxygenation (*44*), and photodynamic priming enhances responses to immune checkpoint blockade through immunological mechanisms involving DAMP release and dendritic-cell activation (*45*). These observations support the idea that the vascular consequences of PDT can extend beyond direct tumor injury. Unlike studies of photodynamic priming, which reduce the PS or light dose, the present comparison changed only the power density. Our findings extend this concept by suggesting that the vascular state during PDT determines not only how the tumor microenvironment is modified but also whether signals generated in the tumor reach the circulation.

Systemic immunity after PDT has mostly been attributed to immunogenic cell death and evaluated using soluble DAMPs, mainly HMGB1, HSP70, and ATP (*10–13*). In our experiments, however, these three markers changed in the same direction and to a similar extent in both treatment groups (**Fig. 7**), although the systemic outcomes differed, and only the proteome of the circulating sEVs clearly distinguished the two conditions (**Fig. 8**). The release and the delivery of DAMPs may therefore be separable events. The appearance of a DAMP in the plasma indicates that cells were killed, whereas the packaging of the DAMP into a vesicle that can be captured by APCs may determine whether the cell death is recognized as an immunological signal. Because most effective treatments kill tumor cells (*46–48*), soluble DAMPs may have a limited ability to discriminate between treatment conditions, and the inconsistent reports on PDT-induced immune responses might be partly explained by a variable that soluble DAMP panels cannot detect. This limitation became apparent in this study because we compared the two conditions matched for the total light dose and analyzed the vesicular fraction together with the soluble one. This issue may not be specific to PDT. Radiotherapy, chemotherapy, and oncolytic virotherapy combined with immunotherapy have also been evaluated with the same soluble markers, and a treatment that releases DAMPs but restricts their dissemination would be judged immunogenic even if it fails to induce a systemic response.

Fluence rate, the term used for power density in earlier studies, has long been recognized as a determinant of local tumor control, and an oxygen-conserving regimen was reported to give higher cure rates than an oxygen-depleting one at the same total fluence (24). However, that study used a different PS, a different drug-to-light interval, and a different tumor model, and the endpoint was the irradiated tumor itself. Whether the power density also governs the systemic response has not been examined, and the circulating sEVs, in which the two conditions differed most clearly in this study, have not been analyzed in this context. The photochemical reaction was also evaluated differently. Previous studies on the fluence rate effect estimated the photochemical situation from surrogate measures, either electrode measurements of the tissue oxygen partial pressure (pO_2_) or immunohistochemical hypoxia markers (19, 25), whereas we detected ^1^O_2_ directly in living tumors (**Fig. 3**). This allowed us to follow how ^1^O_2_ generation and the balance between oxygen consumption and supply changed with power density. These findings are also relevant to the clinical use of PDT. Clinical protocols specify the PS dose, the wavelength, and the total fluence, whereas the power density is generally set high so that a session can be completed quickly. Thus, the power density, which is generally chosen for the convenience of the treatment schedule and is frequently neither reported nor matched between studies, may govern the systemic outcome of PDT. Lowering the power density requires no new PS, no change in total light dose, and no new hardware, but clinical irradiation time is limited; delivering 50 J/cm^2^ at 12.5 mW/cm^2^ takes more than 1 h. Irradiation conditions that maintain the balance between oxygen consumption and supply within an acceptable session time must therefore be determined, which requires monitoring of the photochemical reaction during treatment. Measurement of ^1^O_2_ phosphorescence should be one of the most promising approaches (**Fig. 3**). When the session time does not allow a low power density, the increased PD-L1 expression in unirradiated tumors after HPD-PDT (**Fig. 6G**) suggests that combination with PD-L1 blockade may be beneficial. Meanwhile, daylight PDT, in which lesions are exposed to sunlight for a period of hours, is a form of low-power-density irradiation in which the long exposure compensates for the low power density (*49*). A systematic analysis of immune response after daylight PDT may also be informative for better understanding of the abscopal effects.

This study has some limitations. Blocking the vascular response or depleting the sEVs will be required to establish that the sEVs are necessary for the abscopal effect. Because the sEVs were isolated from blood, the contributions of tumor cells, vascular endothelial cells, and infiltrating immune cells could not be distinguished, and the roles of many differentially abundant sEV proteins in antitumor immunity remain to be determined. In addition, we examined one tumor model and one PS, and the photophysical measurements used pulsed excitation at power densities different from those used for treatment. Defining the quantitative relationship between photochemical stress and sEV cargo, including the point at which the cargo shifts from tissue-injury-associated signals toward acute-phase and coagulation-associated proteins, will more clearly elucidate how irradiation conditions shape the systemic response. Extending these analyses to other tumor types and investigating whether circulating sEV profiles correlate with treatment response in patients will be the focus of future studies. Overall, our findings show that power density, a parameter usually chosen for convenience, governs both the local and systemic outcomes of PDT.

## MATERIALS AND METHODS

### Animal model and PDT conditions

Four-week-old female BALB/c mice (Japan SLC, Inc., Shizuoka, Japan) were purchased and acclimated to the laboratory environment for one week prior to experiments. CT26 murine colon carcinoma cells were subcutaneously inoculated into the bilateral dorsal flanks of mice at a density of 1×10^5^ cells per site. PDT was performed 14 days after inoculation. For PDT, mice received an intravenous injection of talaporfin sodium via the tail vein at a dose of 10 mg/kg. Four hours after administration, mice were anesthetized with isoflurane, and the left-side tumor was irradiated using a 680 nm laser. The light source was positioned approximately 2 cm above the tumor surface, generating a circular irradiation spot with a diameter of 1 cm, sufficient to cover the entire tumor area on one side. Mice in the control group received talaporfin sodium injection but were not subjected to laser irradiation. Tumors were irradiated at five power densities (200, 100, 50, 25, and 12.5 mW/cm^2^), and the irradiation time was adjusted to deliver 50 J/cm^2^ in all groups (**Table 1)**. After irradiation, mice were returned to their cages and maintained for an additional 14 days. Body weight and tumor size on both sides were measured daily at a fixed time point. Tumor volume was calculated as length × width^2^/2. Mice were euthanized on day 14 following PDT treatment. All animal procedures were performed in accordance with institutional guidelines and were approved by the Animal Care and Use Committee of The University of Tokyo (approval numbers: A2023AS004-03 and A2026AS007).

### In vivo ^1^O_2_ detection

Female BALB/c mice (4 weeks old) were used. All animal procedures were performed in accordance with the guidelines described above. When the tumor length reached approximately 8 mm, talaporfin sodium was administered via tail vein injection at a dose of 10 mg/kg. Data acquisition was started 4 h after administration. Time-resolved *in vivo* ^1^O_2_ phosphorescence was detected using a near-infrared photomultiplier tube (PMT)-based detection system (TCMPC1270, SHB Analytics, Berlin, Germany). Two band-pass filters with a full width at half maximum of approximately 40 nm were used to discriminate ^1^O_2_ phosphorescence from background emission. The first filter had maximum transmission at 1270 nm and collected ^1^O_2_ phosphorescence together with background signals such as PS phosphorescence and short-term artifacts. The second filter had maximum transmission at 1200 nm and collected background emission with negligible contribution from ^1^O_2_ phosphorescence. The ^1^O_2_ signal was obtained by subtracting the scaled 1200 nm signal from the 1270 nm signal. Although the fiber position was fixed, the prolonged duration of the measurements meant that small movements of the mouse, as well as other factors such as PS photobleaching, could affect the signal intensity. Moreover, the PS phosphorescence intensity at the two wavelengths was not necessarily identical. Because these factors were expected to affect the amplitude rather than the decay kinetics, a scaling factor close to 1 was applied to the 1200 nm signal before subtraction. The resulting difference signals were fitted with a standard biexponential function.

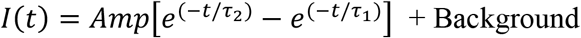

where *I*(*t*) represents the detected ^1^O_2_ phosphorescence counts at time *t* after the excitation pulse. *Amp* is an amplitude parameter proportional to the amount of ^1^O_2_ generated. *τ*_1_corresponds to the lifetime of ^1^O_2_, whereas *τ*_2_ represents the apparent lifetime of the triplet-state PS. Under low-oxygen conditions *in vivo*, *τ*_1_ is smaller than *τ*_2_ (*50*). Different average power densities were achieved by varying the pulse repetition frequency.

### Cell culture

CT26 murine colon carcinoma cells were used throughout. CT26.WT cells (ATCC CRL-2638) were obtained from the American Type Culture Collection (ATCC, Manassas, VA, USA). Cells were cultured in RPMI-1640 medium supplemented with 10% fetal bovine serum (FBS) and 1% penicillin–streptomycin and maintained at 37 °C in a humidified atmosphere containing 5% CO₂. The culture medium was replaced every 2 days, and cells were routinely passaged before reaching full confluence. Cells were used within passages 14–20 for all experiments.

### In vitro photobleaching of the PS

An aqueous solution of talaporfin sodium (10 μM) was irradiated with a halogen lamp at 100 or 25 mW/cm^2^, and fluorescence spectra were recorded between 600 and 800 nm as a function of the delivered light dose.

### In vivo fluorescence imaging of the PS

*In vivo* fluorescence imaging was performed to monitor the photobleaching of talaporfin sodium during PDT. Fluorescence signals were acquired using a Luminous Quester NX imaging system (Shimadzu Corporation, Kyoto, Japan). Mice bearing CT26 tumors received talaporfin sodium (10 mg/kg) by intravenous administration via the tail vein. After a drug-to-light interval of 4 h, fluorescence images were continuously acquired during irradiation of the tumor at 680 nm under the HPD and LPD conditions. Regions of interest (ROIs) were defined over the directly irradiated tumor area and an adjacent non-irradiated reference region. Fluorescence intensity was quantified using the instrument analysis software. Because fluorescence signals were also detected in the non-irradiated reference region after the laser was turned on, the reference-region signal was used to account for non-local fluorescence contributions rather than being treated as a static background. The corrected fluorescence intensity of the irradiated tumor was calculated by subtracting the fluorescence intensity of the reference region from that of the tumor region. The corrected values were subsequently normalized to the corresponding pre-irradiation value and expressed as a percentage of the baseline fluorescence. Changes in normalized talaporfin sodium fluorescence were plotted against the delivered light dose to evaluate and compare the photobleaching kinetics under HPD and LPD irradiation.

### In vivo tumor blood perfusion imaging

Tumor blood perfusion was monitored using an RFLSI-ZW laser speckle blood-flow imaging system (RWD Life Science, Shenzhen, China). This system measures tissue blood flow non-invasively by analyzing dynamic changes in laser speckle patterns generated by coherent laser illumination of the tissue surface. Mice were anesthetized and positioned on the imaging platform, and the tumor region was exposed for imaging. Baseline blood flow images were acquired prior to irradiation. Subsequent measurements were performed after PDT under both HPD and LPD irradiation conditions. Blood perfusion maps were generated in real time, and ROIs were defined over the tumor area for quantitative analysis of relative blood flow changes. To minimize variations in signal intensity caused by differences in animal positioning and imaging distance, an additional ROI was defined in adjacent normal tissue. Tumor blood flow was normalized to the perfusion measured in the neighboring normal tissue, and relative changes before and after PDT were calculated.

### Cytokine analysis in serum and tumor tissue

Blood was collected 6 h after PDT, allowed to clot at room temperature, and centrifuged to obtain serum. Concentrations of IL-2, IL-6, TNF-α, and IFN-γ were quantified using commercially available ELISA kits from Thermo Fisher Scientific according to the manufacturer’s instructions. To evaluate cytokine levels in the unirradiated contralateral tumor, tumors were harvested 14 days after PDT and mechanically dissociated. Briefly, tumors were minced and homogenized, followed by filtration through a 70 μm cell strainer to obtain tumor extracts. Total protein concentration in the tumor lysates was determined by the bicinchoninic acid (BCA) assay, and cytokine concentrations were also measured using the same ELISA kits as mentioned before. Cytokine levels in tumor samples were normalized to the total protein content to allow comparison between samples.

### Tumor dissociation and flow cytometry analysis

Tumors were excised from mice and mechanically minced into small fragments (<0.75 mm in diameter) after removal of surrounding fat, necrotic, and fibrotic tissues. Tumor tissues were enzymatically dissociated using a tumor dissociation kit (Miltenyi Biotec, catalog no. 130-096-730). Briefly, tumor fragments were incubated in RPMI-1640 containing enzyme mixture at 37 °C for 40 min with gentle agitation. The digested tissue suspension was filtered through a 70 μm cell strainer to obtain a single-cell suspension. Red blood cells were lysed with a lysis buffer on ice for 5 min, and the reaction was stopped with PBS. Cells were then centrifuged at 300g for 5 min, resuspended in Cell Staining Buffer, and counted.

For flow cytometric analysis, 1 × 10^6^ cells were aliquoted for each staining condition and analyzed using a CytoFLEX flow cytometer (Beckman Coulter). Fc receptors were blocked using TruStain FcX™ PLUS (BioLegend) on ice for 10 min. Cells were then stained with fluorophore-conjugated antibodies against CD45 and PD-L1 to evaluate tumor cell PD-L1 expression; F4/80, MHC II, and CD206 to characterize tumor-infiltrating macrophage phenotypes; CD3, CD4, and CD8 to analyze infiltrating T cell populations. The antibodies are listed in table **S2**. Antibody staining was performed in the dark on ice for 20 min, followed by washing with Cell Staining Buffer (PBS containing 3% FBS and 2.5 mM EDTA). Dead cells were excluded using 7-aminoactinomycin D (7-AAD) staining prior to acquisition. Samples were analyzed on a flow cytometer and data were processed using FlowJo software (FlowJo, LLC, Ashland, OR, USA). The gating hierarchy is shown in fig. **S4.**

### Measurement of circulating DAMPs

To evaluate the release of DAMPs following PDT, blood samples were collected at 0.5, 6, and 24 h after irradiation. Whole blood was allowed to clot at room temperature for 15 min, followed by incubation at 4°C for 20 min. Samples were then centrifuged at 1,500 g for 15 min at 4°C, and the upper serum layer was transferred to fresh tubes. Serum ATP levels were measured using the ATP Assay Kit-Luminescence (Dojindo Laboratories, Kumamoto, Japan; Cat. No. A550), according to the manufacturer’s instructions. Serum HSP70 and HMGB1 levels were quantified using ELISA kits (HSP70 ELISA Kit, Abcam; HMGB1 ELISA Kit, Proteintech, Cat. No. KE00343), following the manufacturers’ instructions. DAMPs concentrations were determined using the corresponding standard curves.

### Bone marrow-derived macrophage (BMDM) polarization assay

Bone marrow-derived macrophages (BMDMs) were generated from bone marrow cells of 6-week-old BALB/c mice by culturing in RPMI-1640 medium supplemented with 10% FBS, 1% penicillin-streptomycin, and macrophage colony-stimulating factor (M-CSF, 20 ng/mL) for 7 days. CT26 cells were incubated with 10 μM talaporfin sodium for 4 h before halogen-lamp irradiation at 100 mW/cm² for 600 s or 20 mW/cm² for 3000 s (60 J/cm² in both groups). One hour after PDT, culture supernatants were collected and centrifuged (2,000g, 10 min) to remove cellular debris. BMDMs were then incubated with the resulting CM for 48 h, followed by RNA extraction and quantitative real-time polymerase chain reaction (qPCR) analysis of M1-associated markers (CD86, TNF-α, and Irf5) and M2-associated markers (CD206, Ym1, and PPAR-γ). Gene expression levels were normalized to GAPDH. qPCR was performed using GeneAce SYBR™ qPCR Mix α (Nippon Gene Co., Ltd., Tokyo, Japan; cat. no. 319-07683) on a CFX Duet Real-Time PCR System (Bio-Rad). The primer sequences used for qPCR are listed in **table S3**.

### Isolation and nanoparticle tracking analysis of plasma sEVs

Six hours after PDT, blood samples were collected via retro-orbital bleeding into EDTA-coated tubes. Plasma was separated by centrifugation, and sEVs were isolated using the MagCapture Exosome Isolation Kit PS (FUJIFILM Wako Pure Chemical Corporation, catalog no. 290-84103) according to the manufacturer’s instructions. The isolated sEV preparations were diluted fivefold before analysis. Particle concentration and size distribution were determined based on Brownian motion using a NanoSight NS300 instrument (Malvern Panalytical, Tokyo, Japan). Three 60 s recordings were acquired for each replicate, and the resulting measurements were averaged to determine the particle concentration and modal particle size.

### Protein digestion and peptide purification

Proteins in dried sEV samples were digested by the phase-transfer surfactant method as described previously (*51*). Proteins were reduced with 10 mM dithiothreitol (DTT; FUJIFILM Wako Pure Chemical Corporation, Osaka, Japan) for 30 min at 37 °C and subsequently alkylated with 50 mM 2-iodoacetamide (IAA; FUJIFILM Wako Pure Chemical Corporation) for 30 min at room temperature in the dark. The samples were then diluted fivefold with 50 mM ammonium bicarbonate and digested overnight at 37 °C with shaking using 1 μg each of lysyl endopeptidase (Lys-C; FUJIFILM Wako Pure Chemical Corporation) and trypsin (Promega, Tokyo, Japan). The digestion was terminated by adding trifluoroacetic acid (TFA) to a final concentration of 0.5%. The acidified peptide samples were purified using tandem StageTips containing stacked styrene-divinylbenzene (SDB-XC; upper) and strong cation exchange (SCX; lower) Empore disk membranes (GL Sciences, Tokyo, Japan). The StageTips were sequentially washed with 0.1% TFA containing 5% acetonitrile (ACN) and 0.1% TFA containing 80% ACN. Peptides were then sequentially eluted from the SCX membrane with a buffer containing 4% TFA, 30% ACN, and 500 mM ammonium acetate, followed by a buffer containing 30% ACN and 500 mM ammonium acetate. Following elution, the peptide-containing solutions were diluted fivefold with 1% TFA. The peptides were subsequently desalted using StageTips containing SDB-XC Empore disk membranes and reconstituted in loading buffer containing 0.5% TFA and 5% ACN for LC-MS/MS analysis. The concentration of the final peptide preparations was determined by measuring absorbance at 205 nm using a NanoDrop spectrophotometer (Thermo Fisher Scientific) and a mass extinction coefficient of 31 mL mg^-1^ cm^-1^, as described previously (*52*).

### NanoLC-MS/MS analysis

LC-MS/MS analysis was performed in data-dependent acquisition mode using an Orbitrap Exploris 480 mass spectrometer equipped with a high-field asymmetric waveform ion mobility spectrometry (FAIMS) Pro Duo interface (Thermo Fisher Scientific). Nanoelectrospray ionization was performed in positive-ion mode with a spray voltage of 2.4 kV and an ion transfer tube temperature of 275 °C. FAIMS was operated at standard resolution with a total carrier gas flow of 4.0 L/min. The compensation voltages (CVs) were set to −40 and −60 V, and the cycle time for each CV experiment was set to 1 s.

Full MS scans were acquired over an m/z range of 375–1,500 at an Orbitrap resolution of 60,000. The RF lens was set to 50%, the normalized automatic gain control (AGC) target was set to 300% (absolute AGC target, 3.0 × 10^6^), and the maximum injection time was set to automatic. Precursor ions with charge states of 2–7 and intensities above 5.0 × 10^3^ were selected for MS/MS analysis. Dynamic exclusion was applied after a single selection for 20 s with a mass tolerance of ±5 ppm, and isotope exclusion was enabled. Precursor ions were isolated using an isolation window of 1.6 m/z and fragmented by higher-energy collisional dissociation (HCD) with a normalized collision energy of 30%. MS/MS spectra were acquired in the Orbitrap at a resolution of 15,000, with the first mass set to m/z 120, a standard AGC target, and an automatic maximum injection time. Full MS spectra were recorded in profile mode, whereas MS/MS spectra were recorded in centroid mode.

LC separation was performed using an Ultimate 3000 nanoLC system (Thermo Fisher Scientific) equipped with a 250-mm-long, 100-μm-inner-diameter analytical column packed with Reprosil-Pur 120 C18-AQ 1.9-μm reversed-phase material (Dr. Maisch, Ammerbuch, Germany (*53*)). Mobile phase A consisted of 4% ACN and 0.5% acetic acid in water, whereas mobile phase B consisted of 80% ACN and 0.5% acetic acid in water. The autosampler temperature was maintained at 10 °C. Peptide samples were injected at 50 ng per LC-MS/MS run, and the flow rate was set to 400 nL/min. Separation was performed using the following multistep gradient: 5% B from 0 to 5 min; 5–19% B from 5 to 60.3 min; 19–29% B from 60.3 to 81.3 min; 29–40% B from 81.3 to 90.0 min; and 40–99% B from 90.0 to 90.1 min. The column was washed at 99% B from 90.1 to 95.0 min, returned to 5% B from 95.0 to 95.1 min, and re-equilibrated at 5% B until 130 min.

### Database searching and proteomic data analysis

Raw MS data were analyzed using MaxQuant version 2.6.7.0 (*54*). Peptides and proteins were identified by searching with Andromeda against the Mus musculus Swiss-Prot database (May 2026 release; 17,274 protein entries). Raw files acquired using FAIMS were split into MaxQuant-compatible, CV-specific mzXML files using the FAIMS MzXML Generator (https://github.com/coongroup/FAIMS-MzXML-Generator). The precursor mass tolerances were set to 20 ppm for the first search and 4.5 ppm for the main search, and the fragment ion mass tolerance was set to 20 ppm. Trypsin/P was specified as the proteolytic enzyme, allowing up to two missed cleavages. Carbamidomethylation of cysteine was specified as a fixed modification, whereas oxidation of methionine and acetylation of the protein *N*-terminus were specified as variable modifications. Peptide-spectrum matches (PSMs) and protein identifications were filtered at a false discovery rate (FDR) of <1%. Match between runs (MBR) was enabled using the default settings, with a match time window of 0.7 min and an alignment time window of 20 min. Protein groups marked as “Only identified by site,” “Potential contaminant,” or “Reverse” were excluded from subsequent analyses. Protein intensities were log_2_-transformed and median-normalized. Proteins quantified in at least two of three biological replicates in at least one group were retained. Differences between two groups were tested with a two-sided [specify test], and proteins with a fold change ≥ 2 and *P* < 0.05 were considered differentially expressed. Spearman correlation coefficients were used for hierarchical clustering of the samples.

### Statistical analysis

Data are presented as mean ± standard deviation (SD), unless otherwise indicated. Tumor growth curves are presented as mean ± standard error of the mean (SEM). Comparisons among three or more groups were performed using one-way analysis of variance (one-way ANOVA) followed by Tukey’s multiple comparisons test. Tumor growth curves were analyzed using two-way repeated-measures ANOVA followed by Tukey’s multiple comparisons test. Statistical analyses were performed using GraphPad Prism 10 (GraphPad Software, San Diego, CA, USA). Differences were considered statistically significant at *P* < 0.05. The number of animals or samples in each experiment is given in the figure legends.

## Supporting information

Proteomics data

## Funding

Japan Science and Technology Agency FOREST Program JPMJFR215E (T.N.)

Japan Science and Technology Agency Program for co-creating startup ecosystem JPMJSF2313 (T.N.)

Japan Society for the Promotion of Science JP25K02525, JP25K22571, JPJSBP120253503 (T.N.)

Nakatani Foundation (T.N.)

Japan Agency for Medical Research and Development PRIME program JP24gm7010003 (E.K.)

Japan Science and Technology Agency ASTEP JPMJTR25U9 (E.K.)

Japan Society for the Promotion of Science JP26K22909, JP26K02156 (E.K.)

## Author contributions

Conceptualization: H.J., T.N.

Methodology: H.J., E.K., S.W., S.H., T.N.

Investigation: H.J., E.K., S.W., S.H.

Visualization: H.J., E.K.

Supervision: T.N.

Writing—original draft: H.J.

Writing—review & editing: E.K., S.W., S.H., T.N.

## Competing interests

S.H. is the co-founder of SHB Analytics.

The mass spectrometry proteomics data have been deposited in the ProteomeXchange Consortium via the jPOST partner repository under the dataset identifier PXD 086974 (JPST005066). This study did not generate new materials or custom code.

