## Supplementary Figures and Tables for "Light power density governs local and systemic responses to photodynamic therapy"

Hanqing Jiang *et al.*

**This PDF file includes:**

Figs. S1 to S4
Tables S1 and S2

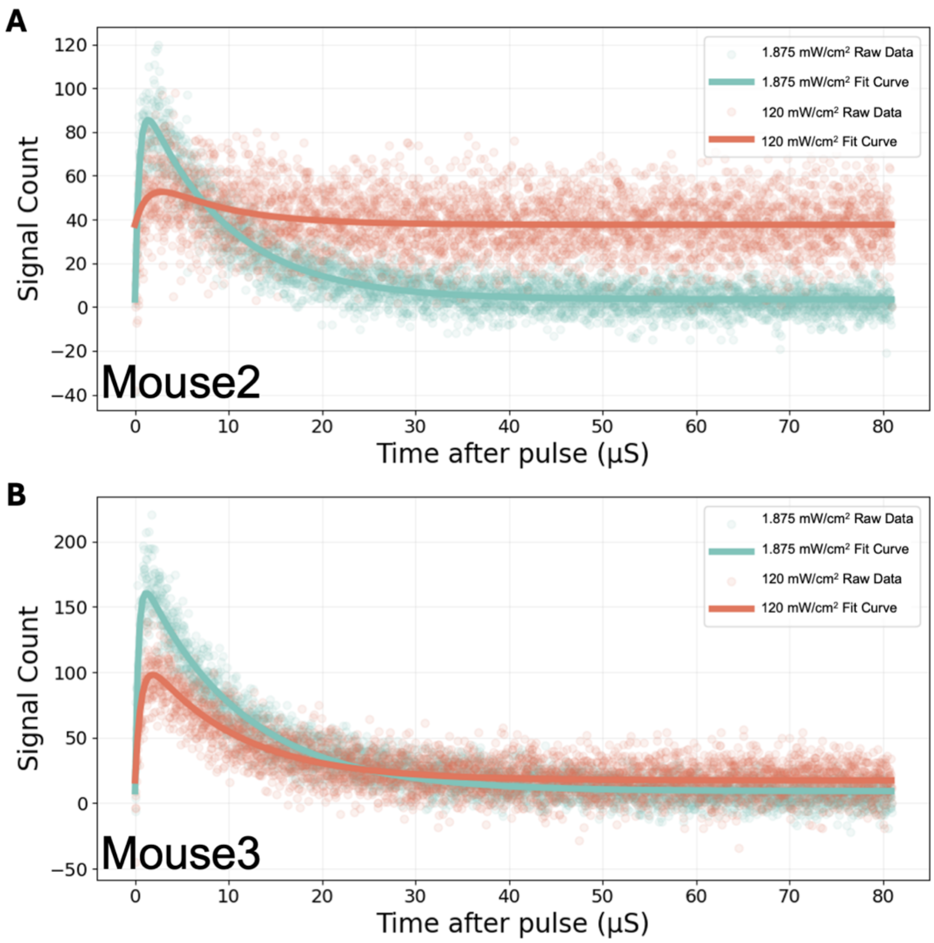


**Fig. S1. Additional representative *in vivo* singlet oxygen luminescence decay curves from independent mice.** (A, B) $^1O_2$ luminescence decay curves obtained from the remaining two independent mice under different irradiation power densities. Experimental data were fitted using a biexponential decay model. Curves of different colors represent measurements obtained under different laser pulse parameters. Spheres: experimental points; line: curve fitting.

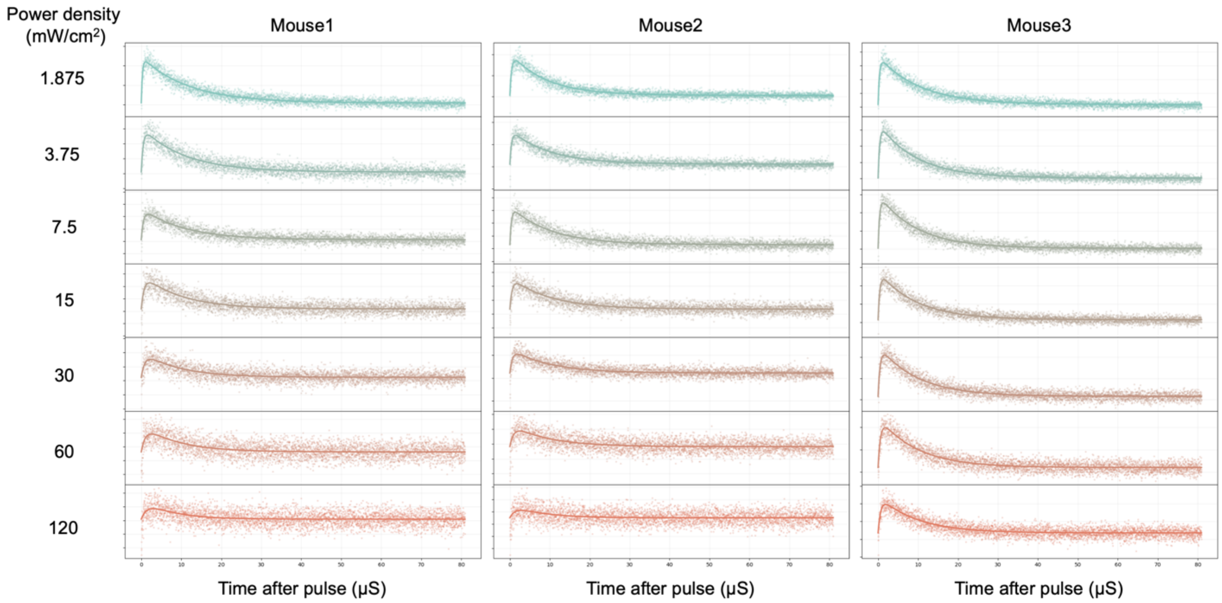


**Fig. S2. Time-resolved $^1O_2$ phosphorescence decay curves and biexponential fitting across irradiation power densities.** Time-resolved $^1O_2$ phosphorescence signals were acquired from three mice under seven irradiation power densities ranging from 1.875 to 120 mW/cm$^2$. Columns represent individual mice, and rows represent the corresponding irradiation power densities. Spheres indicate experimentally measured data points, and solid lines indicate the corresponding biexponential fits. The complete set of decay curves illustrates the power-density-dependent changes in $^1O_2$ phosphorescence kinetics used to derive the amplitude (*Amp*) and characteristic decay parameters $\tau_1$ and $\tau_2$ presented in **Fig. 3**.

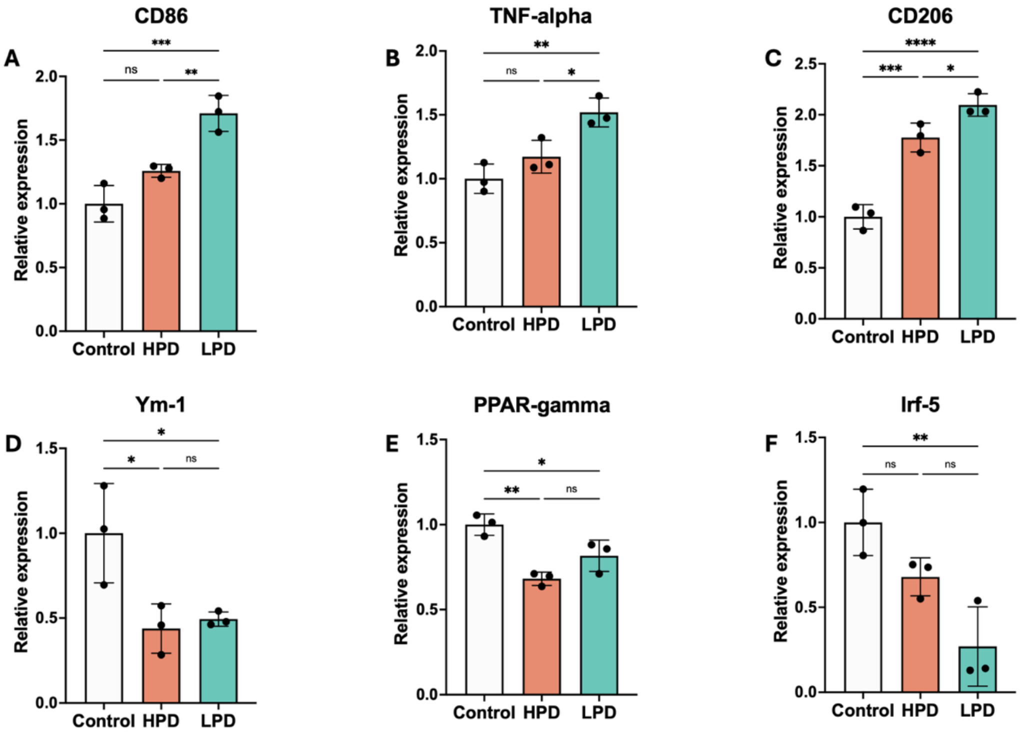


**Fig. S3. Conditioned media from PDT-treated CT26 cells promote macrophage polarization.** Bone marrow-derived macrophages (BMDMs) were cultured with conditioned media collected from CT26 cells following PDT under different irradiation power densities. Relative mRNA expression levels of **(A)** *CD86*, **(B)** *TNF-α*, **(C)** *CD206*, **(D)** *Ym-1*, **(E)** *PPAR-γ*, and **(F)** *Irf-5* were determined by quantitative PCR. Data are presented as mean ± SD.

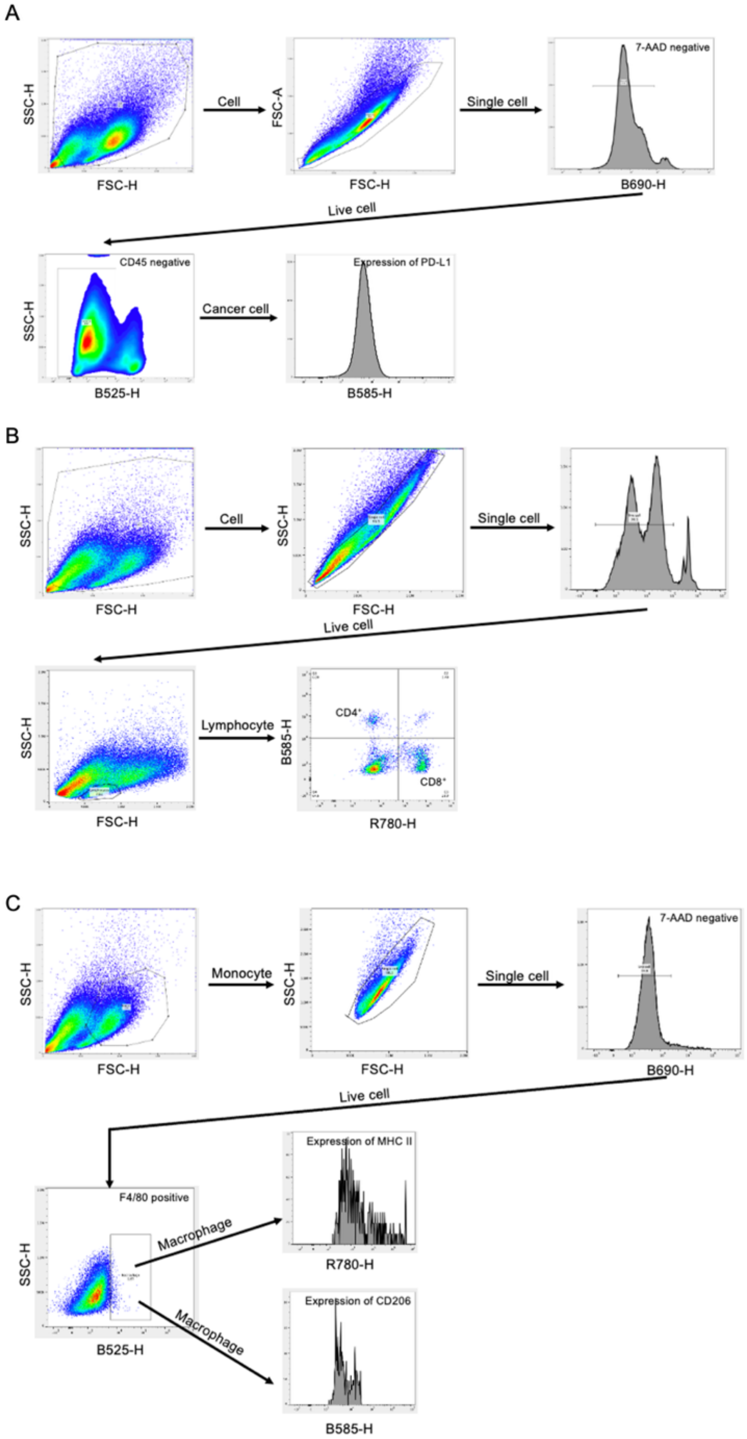


**Fig. S4. Flow cytometric gating strategy for immune profiling of tumor tissues.** (A) Representative gating strategy for identifying CD45$^{-}$ tumor cells and analyzing PD-L1 expression. (B) Representative gating strategy for identifying tumor-infiltrating lymphocytes and determining the proportions of CD4$^{+}$ and CD8$^{+}$ T cells. (C) Representative gating strategy for identifying tumor-infiltrating macrophages and analyzing MHC II and CD206 expression.

**Table S1. Selected sEV proteins and their relative abundance following PDT.**

| Description | Protein name | Mean $log_2$ normalized abundance (Control) | Mean $log_2$ normalized abundance (HPD) | Mean $log_2$ normalized abundance (LPD) |
|---|---|---|---|---|
| Liver-derived acute-phase reactants | SAA1 | 26.24 | 30.06 | 28.41 |
| | SAA2 | 23.90 | 29.82 | 27.95 |
| | CRP | 25.58 | 27.12 | 26.58 |
| Myeloid activation markers | CD14 | 18.08 | 22.97 | 20.59 |
| Endothelial disruption | von Willebrand factor | 19.14 | 21.15 | 19.77 |
| Coagulation cascade factors | Coagulation factor IX | 21.06 | 22.28 | 21.55 |
| | Coagulation factor XI | 19.67 | 20.77 | 20.07 |
| | Coagulation factor XIII | 22.01 | 23.16 | 22.88 |
| | Fibrinogen beta chains | 28.129 | 29.76 | 29.04 |
| | Fibrinogen gamma chains | 28.69 | 30.49 | 29.67 |
| Cell-death-associated proteins | GSDMA2 | 19.63 | 21.71 | 19.19 |
| sEV-phagocytosis bridging molecules | TIM-4 | 27.88 | 26.52 | 27.53 |
| DAMP- and alarmin-associated proteins | H2A | 23.31 | 24.18 | 25.712 |
| | H4 | 22.01 | 22.91 | 24.20 |
| | S100A9 | 23.49 | 23.27 | 24.41 |
| | Lysozyme C-1 | 22.74 | 21.99 | 23.90 |
| Facilitate targeted sEV phagocytosis | Annexin A6 | 26.49 | 25.39 | 26.93 |
| Macrophage functional polarization | FABP5 | 22.19 | 20.86 | 23.36 |
| Immune synapse formation | Plastin-2 | 19.98 | 20.96 | 21.58 |

**Table S2. Antibodies used for flow cytometry.**

| Marker | Fluorophore | Clone | Supplier | Catalog No. |
|---|---|---|---|---|
| CD45 | FITC | S18009F | BioLegend | 157213 |
| PD-L1 | PE | 10F.9G2 | BioLegend | 124307 |
| F4/80 | FITC | BM8 | BioLegend | 123107 |
| MHC II | APC/Cy7 | M5/114.15.2 | BioLegend | 107627 |
| CD206 | PE | C068C2 | BioLegend | 141705 |
| CD3 | FITC | 17A2 | BioLegend | 100203 |
| CD4 | PE | GK1.5 | BioLegend | 100407 |
| CD8 | APC/Cy7 | 53-6.7 | BioLegend | 100713 |
